# Removing zero-lag functional connections can alter EEG-source space networks at rest

**DOI:** 10.1101/617118

**Authors:** Jennifer Rizkallah, Hassan Amoud, Matteo Fraschini, Fabrice Wendling, Mahmoud Hassan

## Abstract

Electro-encephalography (EEG) source connectivity is an emerging approach to estimate brain networks with high time/space resolution. Here, we aim to evaluate the effect of different functional connectivity (FC) methods on the EEG-source space networks at rest. The two main families of FC methods tested are: i) the FC methods that do not remove the zero-lag connectivity including the Phase Locking Value (*PLV*) and the Amplitude Envelope Correlation (*AEC*) and ii) the FC methods that remove the zero-lag connections such as the Phase Lag Index (*PLI*) and orthogonalisation approach combined with *PLV* (*PLV*_*orth*_) and *AEC* (*AEC*_*orth*_). Methods are evaluated on resting state dense-EEG signals recorded from 20 healthy participants. Networks obtained by each FC method are compared with fMRI networks at rest (from the Human Connectome Project -HCP-, N=487). Results show low correlations for all the FC methods, however *PLV* and *AEC* networks are significantly correlated with fMRI networks (ρ = 0.12, *p* = 1.93×10^−8^ and ρ = 0.06, *p* = 0.007, respectively), while other methods are not. These observations are consistent for each EEG frequency bands and for different FC matrices threshold. Furthermore, the effect of electrode density was also tested using four EEG montages (dense-EEG 256 electrodes, 128, 64 and 32 electrodes). Results show no significant differences between the four EEG montages in terms of correlations with the fMRI networks. Our main message here is to be careful when selecting the FC methods and mainly those that remove the zero-lag connections as they can affect the network characteristics. More comparative studies (based on simulation and real data) are still needed in order to make EEG source connectivity a mature technique to address questions in cognitive and clinical neuroscience.

## Introduction

Electro/magneto-encephalography (EEG/MEG) source-space connectivity is a unique non-invasive technique, which enables the tracking of large-scale brain network dynamics on a sub-second time-scale (Hassan and Wendling, 2018; O’Neill et al., 2018; Schoffelen and Gross, 2009). Benefiting from the excellent time resolution of the M/EEG (sub-millisecond), the method consists of identifying brain networks in the cortical space through sensor-level signals. However, several methodological choices should be carefully accounted for to avoid pitfalls.

In this regard, the spatial leakage (presence of spurious connections) was considered as one of the main challenges that affects the accuracy of the M/EEG source-space networks. It was shown to lead to false positive observations caused directly by signal mixing or arising indirectly from the spread of signals from true interacting sources to nearby false loci (Palva et al., 2018; Wang et al., 2018). To deal with this problem, most existing approaches are based on the hypothesis that leakage generates inflated connectivity between estimated sources, which manifests as zero-phase-lag correlations. Thus, these methods dealt with the leakage problem by removing the zero lag connections (Nolte et al., 2004; Stam et al., 2007) or adopting orthogonalization-based approach (Brookes et al., 2012; Hipp et al., 2012).

Here we compare two families of functional connectivity (FC) methods: i) the FC methods that do not remove the zero-lag-phase connectivity including the Phase Locking Value (*PLV*) and the Amplitude Envelope Correlation (*AEC*) and ii) the FC methods that remove the zero-lag connections such as the Phase Lag Index (*PLI*) and orthogonalisation approach combined with *PLV* (*PLV*_*orth*_) and *AEC* (*AEC*_*orth*_). Networks obtained by each method were compared with the networks obtained using fMRI (HCP database, N=487). The impact of the EEG channels density (256, 128, 64 and 32) is also investigated.

## Materials and Methods

### Participants

Dense-EEG recordings (256 channels, EGI, Electrical Geodesic Inc.) were collected from twenty healthy participants (10 women and 10 men; mean age, 23 y). Experiments were performed in accordance with the relevant guidelines and regulations of the National Ethics Committee for the Protection of Persons (CPP), (BrainGraph study, agreement number 2014-A01461-46, promoter: Rennes University Hospital), which approved all the experimental protocol and procedures. All participants in the study provided written informed consents. Participants were asked to relax for 10 minutes with their eyes closed during the acquisition without falling asleep.

### Data acquisition and preprocessing

EEG signals were sampled at 1000 Hz, band-pass filtered within 0.1–45 Hz, and segmented into non-overlapping 40 s long epochs (Chu et al., 2012; Fraschini et al., 2016). Electrodes with poor signal quality (amplitude > 100 µV or < −100 µV) have been identified and interpolated using signals recorded by surrounding electrodes. Segments that have more than 20 electrodes interpolated have been excluded from the analysis. Three clean epochs per subject were then used for source estimation. One subject was excluded from the study due to noisy data.

### Estimation of regional time series

First, the MRI template “Colin27” (Holmes et al., 1998) and EEG channel locations were co-registered using Brainstorm (Tadel et al., 2011). The lead field matrix was then computed for a cortical mesh of 15000 vertices using OpenMEEG (Gramfort et al., 2010). The noise covariance matrix was calculated using a long segment of EEG data at rest, as recommended in (Tadel et al., 2011). An atlas-based approach was used to project EEG signals onto an anatomical framework consisting of 68 cortical regions identified by means of the Desikan-Killiany atlas (Desikan et al., 2006). To reconstruct the regional time series, we used the weighted Minimum Norm Estimate (wMNE), widely used in the context of EEG source localization (Gramfort et al., 2012; Hassan et al., 2015; Hauk, 2004; Kabbara et al., 2017; Rizkallah et al., 2018) and showed higher performance than other algorithms in several comparative studies (Hassan et al., 2014; Hassan et al., 2016). The regional time series were then filtered in the different EEG frequency bands: Delta [0.5-4 Hz], Theta [4-8 Hz], alpha [8-13 Hz], beta [13-30 Hz] and gamma [30-45 Hz]. Results are presented in beta band, in which previous studies have reported its importance in driving large-scale spontaneous neuronal interactions (Brookes et al., 2011; de Pasquale et al., 2012), results for other frequency bands are presented in the supplementary materials. Finally, functional networks were computed using EEG source connectivity method (Hassan et al., 2014; Hassan et al., 2015; Rizkallah et al., 2018; Sakkalis, 2011; Schoffelen and Gross, 2009) by measuring the functional connectivity between the reconstructed regional time series (fig 1).

**Figure 1.**
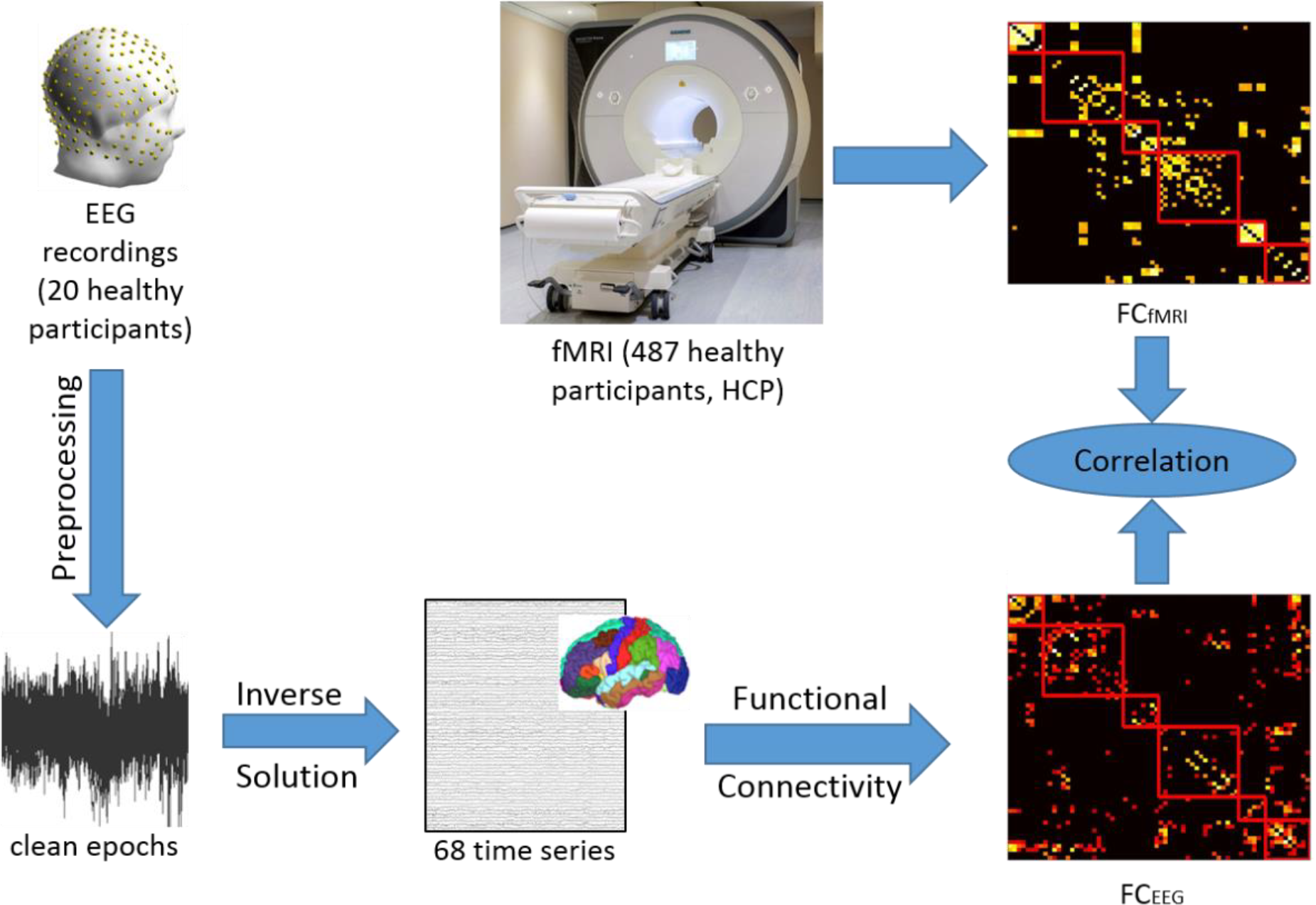
Study pipeline. EEG recordings were preprocessed and clean EEG epochs were used to solve the inverse problem using wMNE. Statistical couplings were then computed between the reconstructed sources using different methods (*PLV*, *AEC*, *PLI*, *PLV*_*orth*_ and *AEC*_*orth*_). Then, the identified matrices were compared with the fMRI functional connectivity matrix obtained from HCP. Abbreviations: EEG: electroencephalogram; wMNE: weighted Minimum Norm Estimate; PLV: phase locking value; AEC: amplitude envelope correlation; PLI: phase lag index; fMRI: functional magnetic resonance imaging; HCP: human connectome project.

### Connectivity measures

The functional connectivity analysis was performed by computing pair-wise statistical interdependence between regional time series using:

#### 1 Phase locking value (PLV)

The phase locking value between two signals *x* and *y* is defined as (Lachaux et al., 1999):

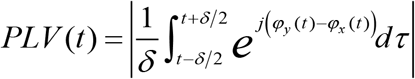

where *φ*_*y*_(*t*) and *φ*_*x*_(*t*) are the phases of the signals *x* and *y* at time *t* extracted using the Hilbert transform. *δ* denotes the size of the window in which *PLV* is calculated. Here, we used a sliding window technique for each epoch to compute the FC matrices. The smallest window length recommended by (Lachaux et al., 2000) was used, equal to 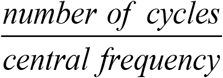 where the number of cycles at the given frequency band is equal to six. Finally, FC were averaged over the 40s epoch.

#### 2 Phase lag index (PLI)

The PLI was introduced as an alternative measure of *PLV* and less sensitive to the influence field spread and amplitude effects. It is defined as follows (Stam et al., 2007):

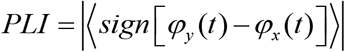

Where *φ*_*y*_(*t*) and *φ*_*x*_(*t*) are the phases of the signals *x* and *y* at time *t* and 〈〉denotes the average over the time.

#### 3 Amplitude envelope correlation (AEC)

The envelopes of the regional time series were estimated using Hilbert transform then Pearson correlation between amplitude envelopes was computed (Brookes et al., 2004).

#### 4 Orthogonalisation approach

The symmetric orthogonalisation approach (Colclough et al., 2015) was used to remove all shared signal at zero lag between regional time series in the time domain. Here, we applied this approach to *PLV* and *AEC* methods.

### fMRI networks

Here, we used data from 487 participants at rest collected from the human connectome project (HCP) (Van Essen et al., 2013). In brief, functional connectivity between each of the 68 regions was assessed by means of analysis of the resting-state fMRI data of the HCP (Q3 release, voxel-size 2 mm isotropic, TR/TE 720/33.1 ms, 1200 volumes, 14:33 minutes). Images were realigned, co-registered with the T1 image, filtered (0.03 - 0.12 Hz), corrected for global effects of motion (realignment parameters), global signal mean, ventricle and white matter signal by means of linear regression and ‘motion-scrubbed’ for potential movement artifacts. Average time-series of the cortical regions were computed by averaging the time-series of the voxels in each of the cortical regions, and functional connectivity between all region pairs was derived by means of correlation analysis. A group-averaged weighted functional connectivity (FC) matrix was formed by averaging the individual matrices, see (van den Heuvel et al., 2016) for more detailed information.

### Statistical comparisons

To statistically assess the difference between the connectivity methods, we thresholded the matrices (EEG and fMRI) by keeping the highest 10% connections (Garrison et al., 2015; Kabbara et al., 2017), results for other threshold values are presented in the supplementary materials. Then, Spearman correlation values between EEG connectivity matrices and the averaged fMRI connectivity matrix were calculated for each participant. Mann-Whitney U Test was used to assess the statistical difference between FC methods. In order to investigate the effect of the number of EEG channels on the identified source-space networks, we spatially subsampled the 256 recordings for each subject and derived the recordings from 128, 64 and 32 channels respectively on which the same steps of preprocessing, source reconstruction and connectivity computation have been applied.

## Results

### Effect of connectivity measures

The FC matrices (averaged over subjects) obtained by each of the FC methods (in beta band) are illustrated in figure 2. These matrices were reordered according to brain lobes. The red module represents the occipital lobe, the green one represents the temporal brain regions, the blue section represents the parietal lobe, the purple module represents the frontal regions, the orange section represents the central lobe and the last module in grey represents the cingulate regions (details are presented in supplementary materials Table1). The averaged FC matrix obtained using fMRI is also illustrated. The visual investigation of these results revealed that matrices obtained from *PLV* and *AEC* connectivity methods are more consistent with the fMRI matrix compared to the other three methods after removing zero lag connections. The latter FC methods connections between brain regions were sparser.

**Figure 2.**
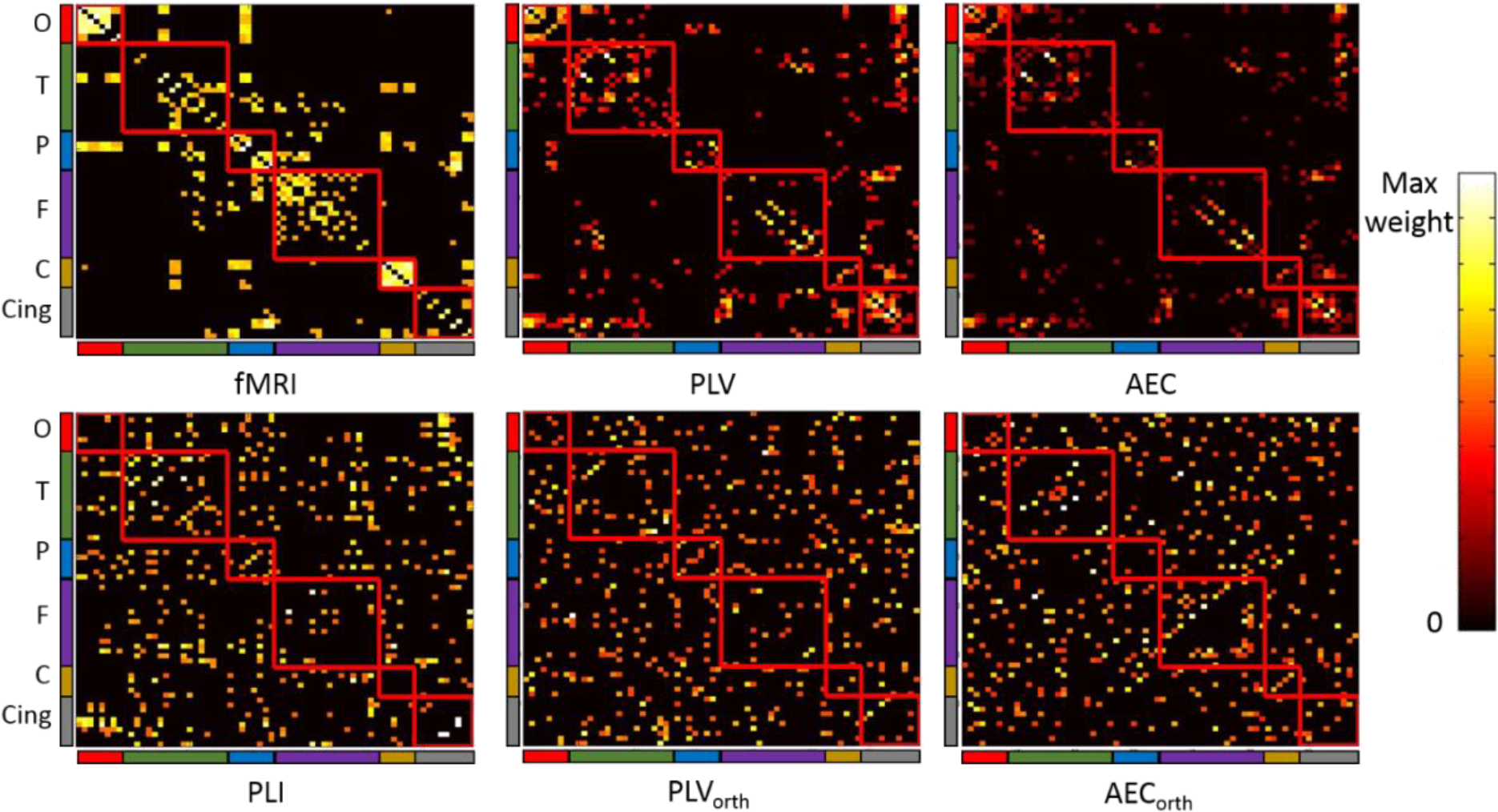
Functional connectivity matrices obtained in beta band from averaged fMRI data and EEG networks. Matrices were ordered according to brain lobes (red: Occipital lobe - O, green: Temporal lobe - T, blue: Parietal lobe - P, purple: Frontal lobe - F, orange: Central lobe - C and grey: Cingulate - Cing). PLV: Phase Locking Value, AEC: Amplitude Envelope Correlation, PLI: Phase Lag Index, PLV_orth_: Phase locking Value after applying leakage correction and AEC_orth_: Amplitude Envelope Correlation after applying leakage correction.

We then explored the Spearman correlations between the EEG network (averaged over subjects) obtained from the five FC methods and the fMRI network at the level of each network connection (edge’s weight) represented in figure 3. Results show low correlations for all the FC methods, however *PLV* and *AEC* networks are significantly correlated with fMRI networks (ρ = 0.12, *p* = 1.93×10^−8^ and ρ = 0.06, *p* = 0.007, respectively). However, the networks obtained after using methods with leakage correction (*PLI*, *PLV*_*orth*_ and *AEC*_*orth*_) were not significantly correlated with fMRI networks (ρ=-0.004, *p*=0.84; ρ=0.03, *p*=0.12 and ρ=0.01, *p*=0.49 respectively).

**Figure 3.**
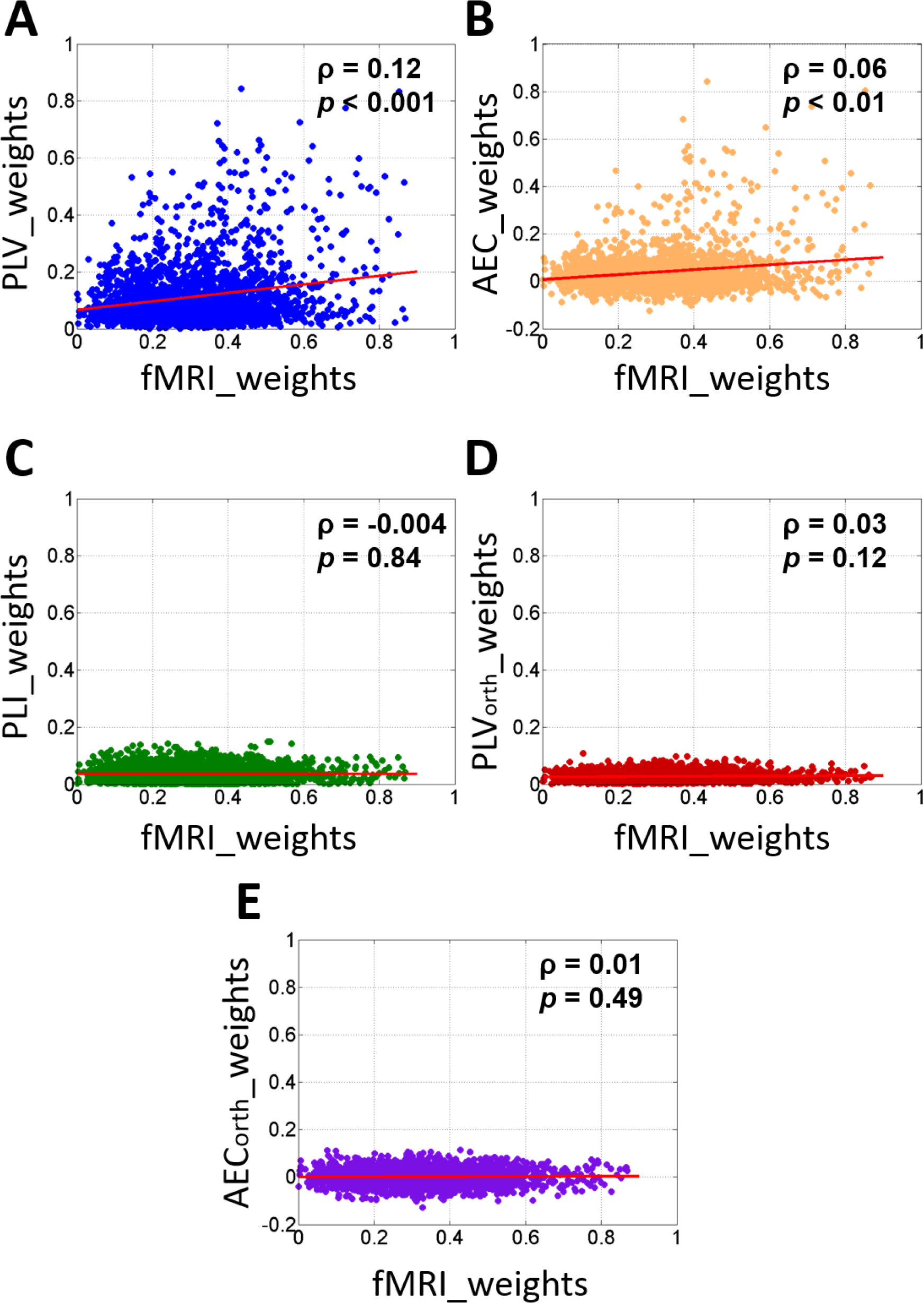
Spearman correlation between different averaged EEG connectivity methods and average fMRI edges weights.

To quantitatively assess the difference between FC methods, Spearman correlation coefficients between FC matrices for each participant and the averaged fMRI connectivity matrix were calculated and presented in figure 4. Results showed significantly higher correlation with fMRI using the *PLV* and *AEC* as compared to the other three methods. *PLV* correlation values were significantly higher than *PLI* (*p*=1.5×10^−7^), *PLV*_*orth*_ (*p*=1.5×10^−7^) and *AEC*_*orth*_ (*p*=5.3×10^−6^). *AEC* correlation values also higher than *PLI* (*p*=2.02×10^−5^), *PLV*_*orth*_ (*p*=2.62×10^−5^) and *AEC*_*orth*_ (*p*=0.001). These results are consistent in the delta, theta, alpha and gamma frequency bands (see figures S1 to S4 in supplementary materials) and after using different thresholds (5%, 20%, 30%, 50% and 80%), see figures S5 to S9 in supplementary materials.

**Figure 4.**
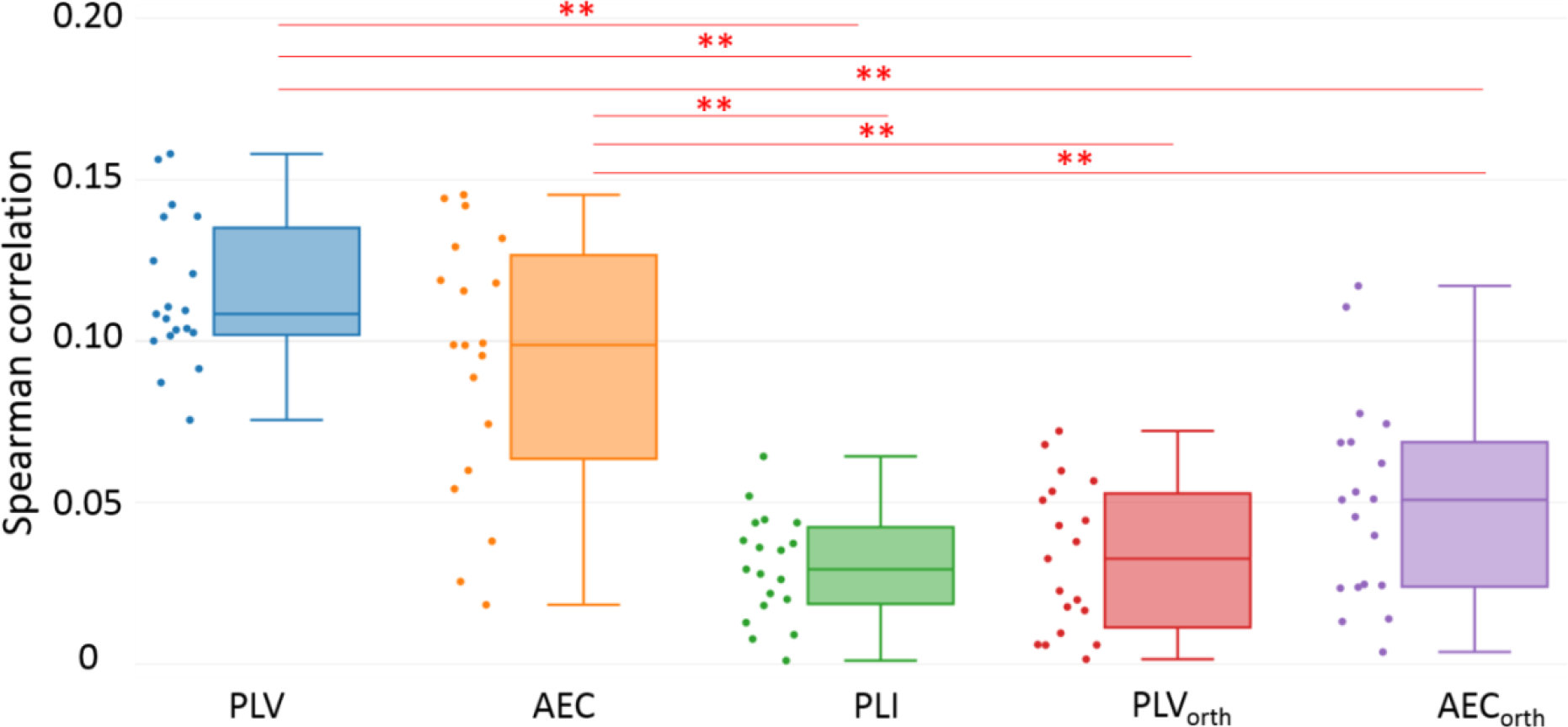
Spearman correlation values between averaged fMRI network and EEG networks in beta band. Individual participant correlations are shown in the scatter plot next to the box plot. ** represents significant differences obtained between methods using Bonferroni correction.

### Effect of electrodes density

In this section, we evaluate the effect of the number of EEG channels on the correlation between EEG and fMRI networks. The FC matrices (averaged over subjects) obtained by *PLV* and *AEC* (the most correlated connectivity methods with fMRI) are illustrated in fig 5.A and fig5.B respectively. The averaged FC matrix obtained using fMRI is also illustrated. The visual investigation of these results revealed that matrices obtained from all EEG montages show similar topologies.

**Figure 5.**
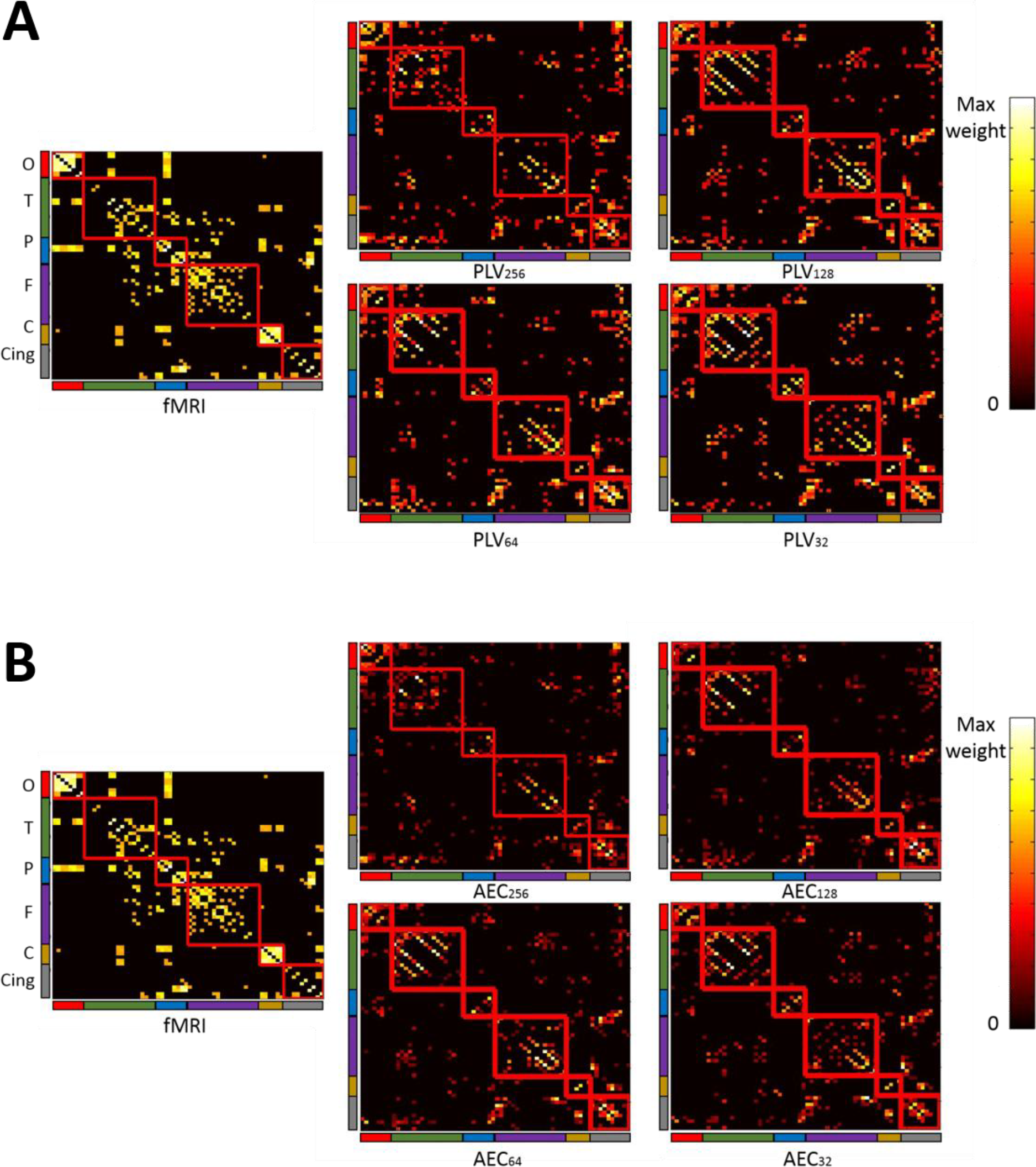
Functional connectivity matrices obtained in beta band from averaged fMRI data and A. PLV and B. AEC networks from four different EEG montages (256, 128, 64 and 32 electrodes). Matrices were ordered according to brain lobes (red: Occipital lobe - O, green: Temporal lobe - T, blue: Parietal lobe - P, purple: Frontal lobe - F, orange: Central lobe - C and grey: Cingulate - Cing).

We Then explored the Spearman correlation (figure 6) between the EEG networks weights obtained using *PLV* and *AEC* methods for each EEG montage (256, 128, 64 and 32 electrodes) and the averaged fMRI network weights. All EEG montages show low but significance correlation with fMRI (*p*<0.001): ρ_*PLV256*_=0.12, ρ_*PLV128*_=0.08, ρ_*PLV64*_=0.11, ρ_*PLV32*_=0.14, ρ_*AEC256*_=0.06, ρ_*AEC128*_=0.09, ρ_*AEC64*_=0.09 and ρ_*AEC32*_=0.14.

**Figure 6.**
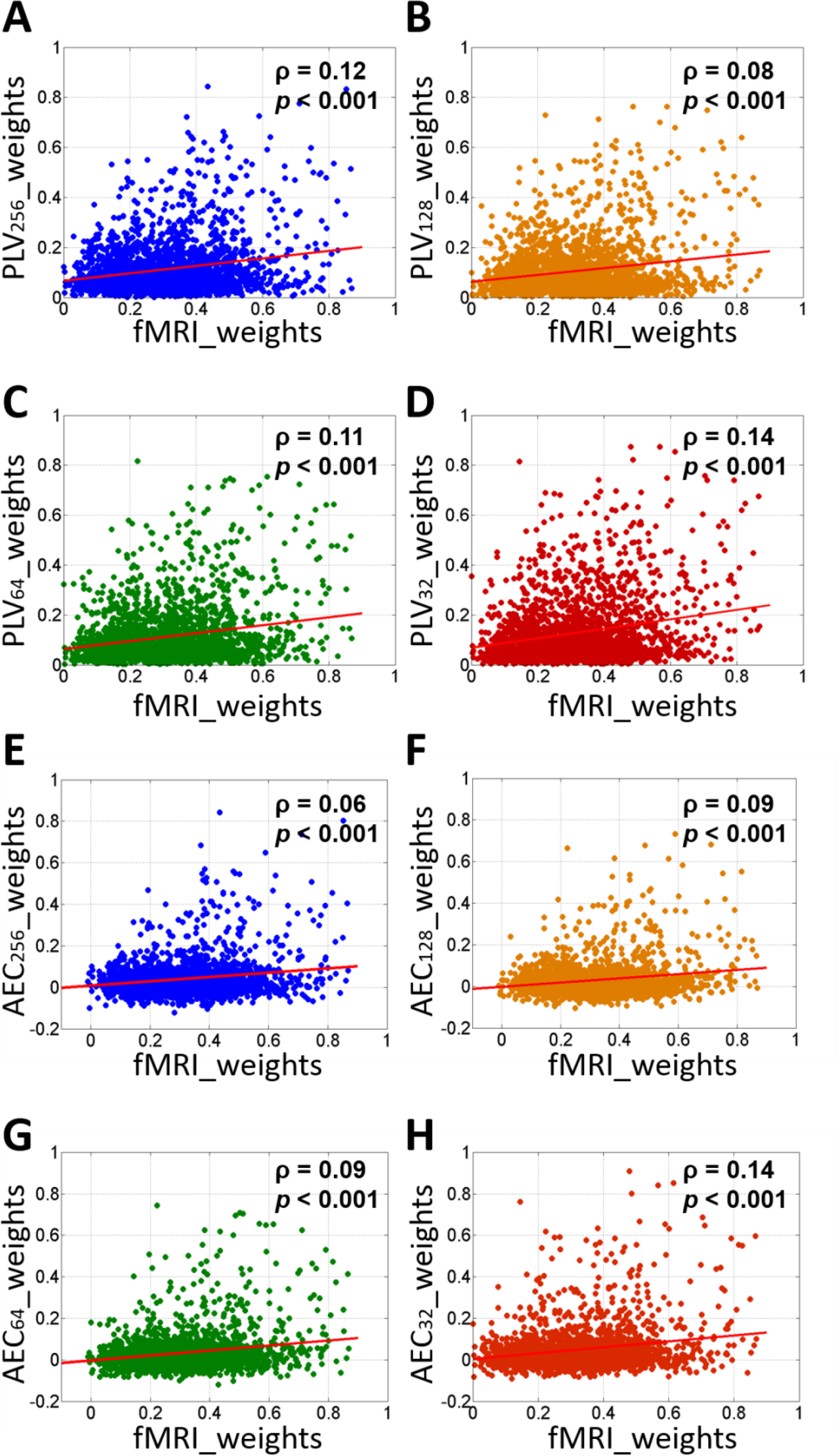
Spearman correlation between fMRI edges weights and different EEG montages using *PLV* (A, B, C and D) and *AEC* (E, F, G and H) methods.

To quantitatively assess the difference between EEG montages, Spearman correlation values between EEG obtained from *PLV* and *AEC* methods and fMRI networks (presented in fig7.A and fig7.B respectively) were computed for each subject for each montage. Comparison between EEG montages was done using Mann-Whitney U Test and shows no significant difference (*p*>0.05). These results are consistent in the delta, theta, alpha and gamma frequency bands (see figures S10 to S13 in supplementary materials) and after using different threshold (5%, 20%, 30%, 50% and 80%), see figures S14 to S18 in supplementary materials.

**Figure 7.**
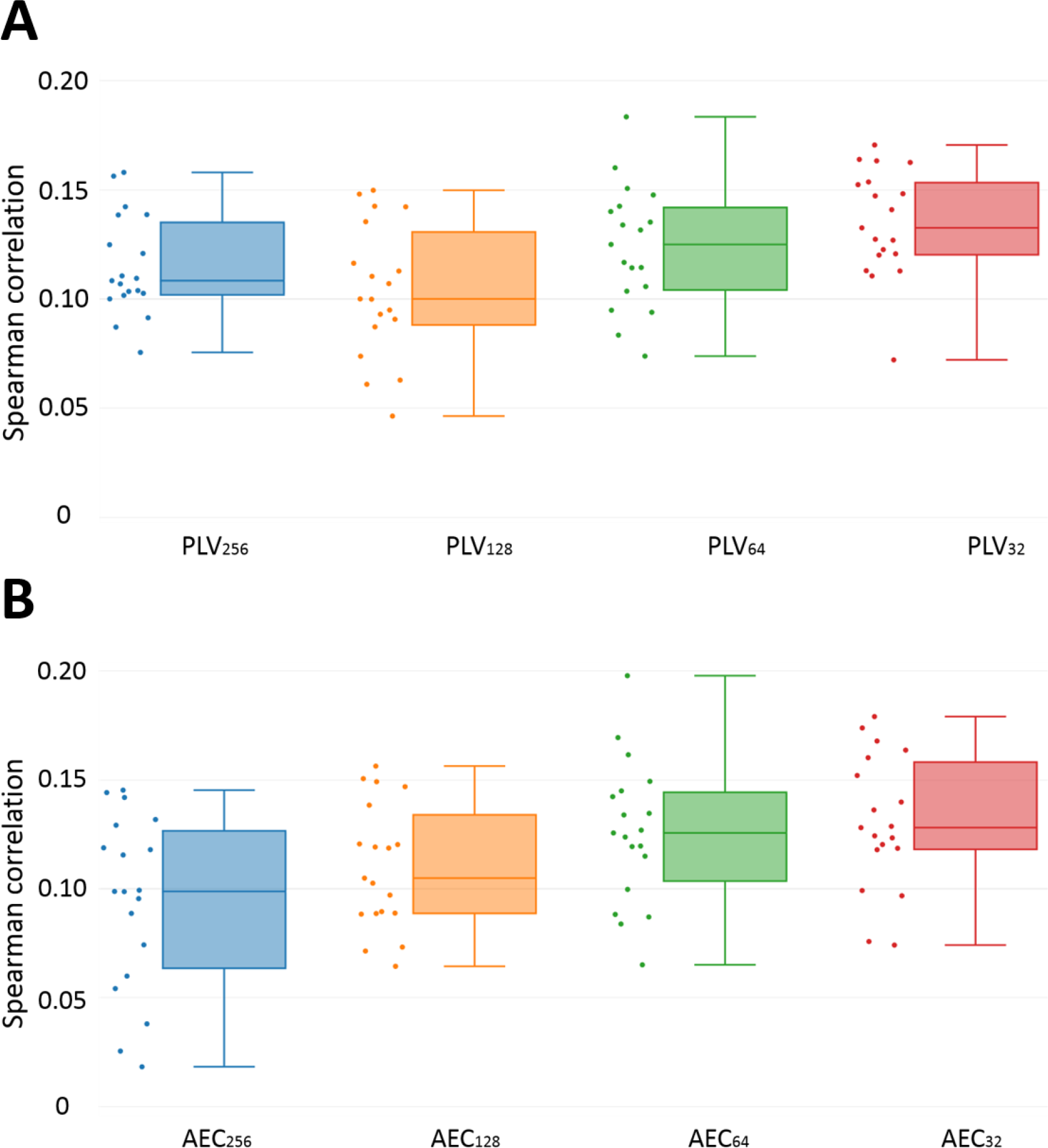
Correlation values between fMRI network and A. *PLV* and B. *AEC* networks in beta band. Individual participant correlations are shown in the scatter plot next to the box plot.

## Discussion

### Connectivity measures

While a large number of FC methods are available, their reliability and consistency are still under exploration. Also, the effect of leakage correction on EEG source-space networks by removing zero lag connections is not sufficiently studied. This paper (and some other recent papers (Colclough et al., 2016)) is a step toward this exploration in which we decided to compare the EEG FC matrices to those obtained using fMRI (HCP databases). Our results show mainly low correlations for all the FC methods. This may be explained by the fact that we are comparing data recorded from different subjects in addition to the intrinsic differences between EEG and fMRI. Slightly higher correlation values between EEG and fMRI resting state networks were found in other study (Liu et al., 2018). Even though the low correlations, our results showed that FC matrices estimated using methods that keep the zero-lag correlations (*PLV* and *AEC)* were significantly correlated with the averaged fMRI functional connectivity matrix as compared to the other methods.

The non-significance between *PLI*, *PLV*_*orth*_ and *AEC*_*orth*_ with fMRI network can be explained by the fact that not all zero-lag connections are spurious. Several previous study described the presence and potential mechanisms for zero-lag connectivity (Gollo et al., 2014; Roelfsema et al., 1997). Recent study showed that removing zero lag connections may indeed reveal false and significantly different estimated connectivity from the true connectivity (Palva et al., 2018). Another study reported that PLV showed the best matching between simulations and empirical data and that zero-lag correlation are very crucial to assess the structural/functional relationships (Finger et al., 2016).

### Electrodes density

Here, we analyzed the effect of EEG electrodes density on the correlation between EEG and fMRI networks. We applied FC methods on dense-EEG (256 electrodes) and the subsampled EEG montages (128, 64 and 32 electrodes). Results showed no significant differences between the different EEG electrodes density. The effect of EEG sensor density on the source localization analysis was previously studied showing that the increase of the EEG channel number may improve the accuracy of the localization (Michel and Murray, 2012; Song et al., 2015). Previous study showed also that Resting State Networks (RSNs) with close brain regions, such as Dorsal Somatomotor Network (DSN), could be successfully reconstructed with low density EEG montage. However, RSNs with distant brain regions, as default mode network (DMN) was affected the most by reducing the number of electrodes (Liu et al., 2018). In addition, previous studies have reported that the results of source localization can be improved with higher densities, but the improvement from 128 to 256 channels is modest (Hassan et al., 2014; Song et al., 2015). We speculate that a compromise between the number of channels and the number of regions of interests should be certainly respected. Our recent findings showed that a high number of electrodes (>32) is mandatory in the case of applications that require higher ‘granularity’, i.e. spatial precision and accurate characterization of the network local properties, such as the identification of epileptogenic networks. Here, and as we were looking to very large-scale properties (global correlation between EEG and fMRI matrices), a low number of electrodes was sufficient. In other context (such as localization of epileptogenic network), it was shown that higher number than standard montage (>32 channels) is needed (Michel and Murray, 2012; Song et al., 2015).

### Methodological considerations

First, in this study the fMRI connectivity matrices were used as a ‘reference’ in order to evaluate the results of each of the FC connectivity measures applied to EEG regional time series. However, the EEG and fMRI data were not collected from the same participants. To that end, we used an averaged matrix over a large number of healthy participants (N=487). We were aware about this limitation and that the ideal situation was to have EEG and fMRI recordings for the same subjects. Of course, the fMRI matrices cannot be considered as an absolute ‘ground truth’ as preprocessing and analysis choices can produce different results (Carp, 2012). However, the spatial resolution of the fMRI networks (not affected by the leakage issue) and the consistency of these networks (we have tested another fMRI-based dataset over 10 healthy subjects and the two networks were highly correlated: correlation is equal to 0.8) can indeed justify its use as performance criteria for analyzing the EEG source-space networks.

Second, the connectivity matrices were thresholded by keeping only the highest 10% connectivity values. It was used to standardize the comparison between the two connectivity methods and fMRI matrices, as network measures are stable across proportional thresholds, as opposed to absolute thresholds (Garrison et al., 2015). We are aware about the effect of this threshold and we have tested other thresholds and the results are very consistent over different threshold values (see figures S5 to S9 and S14 to S18 in supplementary materials).

## Conclusion

M/EEG source connectivity is a unique tool to identify high resolution functional brain networks in time and space. However, results are dependent on the choice of processing methods. In this paper, we analyzed the impact of the method used to measure the functional connectivity and the effect of EEG sensor density. Our results showed that among the different connectivity measures tested, *PLV* and *AEC* provided closer results to fMRI network compared to the three other methods that removes the zero-lag connections. Furthermore, no significant differences were found when using a reduced number of EEG electrodes. We believe that more comparative studies (based on simulation and real data) should be done to make M/EEG source connectivity a mature technique to address questions in cognitive and clinical neuroscience.

## Acknowledgements

This study was supported by the Future Emerging Technologies (H2020-FETOPEN-2014-2015-RIA under agreement No. 686764) as part of the European Union’s Horizon 2020 research and training program 2014–2018. This work has received a French government support granted to the CominLabs excellence laboratory and managed by the National Research Agency in the “Investing for the Future” program under reference ANR-10-LABX-07-01. This work was also financed by the AZM and SAADE Association, Tripoli, Lebanon and by the National Council for Scientific Research (CNRS) in Lebanon. Authors would like to thank Campus France, Programme Hubert Curien CEDRE (PROJET N° 42257YA), for supporting this study. HCP data was provided by the Human Connectome Project, WU-Minn Consortium (Principal Investigators: David Van Essen and Kamil Ugurbil; 1U54MH091657) funded by the 16 NIH Institutes and Centers that support the NIH Blueprint for Neuroscience Research; and by the McDonnell Center for Systems Neuroscience at Washington University. Authors would like to thank Olivier Dufor for collecting the EEG data and Martijn Van Den Heuvel for providing the fMRI connectivity matrices from the human connectome project.

## References

Brookes, M.J., et al., 2004. A general linear model for MEG beamformer imaging. NeuroImage. 23, 936–946.

Brookes, M.J., et al., 2011. Investigating the electrophysiological basis of resting state networks using magnetoencephalography. Proceedings of the National Academy of Sciences. 201112685.

Brookes, M.J., Woolrich, M.W., Barnes, G.R., 2012. Measuring functional connectivity in MEG: a multivariate approach insensitive to linear source leakage. Neuroimage. 63, 910–920.

Carp, J., 2012. On the plurality of (methodological) worlds: estimating the analytic flexibility of fMRI experiments. Frontiers in neuroscience. 6, 149.

Chu, C.J., et al., 2012. Emergence of stable functional networks in long-term human electroencephalography. Journal of Neuroscience. 32, 2703–2713.

Colclough, G.L., et al., 2015. A symmetric multivariate leakage correction for MEG connectomes. NeuroImage. 117, 439–448.

Colclough, G.L., et al., 2016. How reliable are MEG resting-state connectivity metrics? Neuroimage. 138, 284–293.

de Pasquale, F., et al., 2012. A cortical core for dynamic integration of functional networks in the resting human brain. Neuron. 74, 753–764.

Desikan, R.S., et al., 2006. An automated labeling system for subdividing the human cerebral cortex on MRI scans into gyral based regions of interest. Neuroimage. 31, 968–980.

Finger, H., et al., 2016. Modeling of Large-Scale Functional Brain Networks Based on Structural Connectivity from DTI: Comparison with EEG Derived Phase Coupling Networks and Evaluation of Alternative Methods along the Modeling Path. PLoS Comput Biol. 12, e1005025.

Fraschini, M., et al., 2016. The effect of epoch length on estimated EEG functional connectivity and brain network organisation. Journal of neural engineering. 13, 036015.

Garrison, K.A., et al., 2015. The (in) stability of functional brain network measures across thresholds. Neuroimage. 118, 651–661.

Gollo, L.L., et al., 2014. Mechanisms of zero-lag synchronization in cortical motifs. PLoS computational biology. 10, e1003548.

Gramfort, A., et al., 2010. OpenMEEG: opensource software for quasistatic bioelectromagnetics. Biomedical engineering online. 9, 45.

Gramfort, A., Kowalski, M., Hämäläinen, M., 2012. Mixed-norm estimates for the M/EEG inverse problem using accelerated gradient methods. Physics in Medicine & Biology. 57, 1937.

Hassan, M., et al., 2014. EEG source connectivity analysis: from dense array recordings to brain networks. PloS one. 9, e105041.

Hassan, M., et al., 2015. Dynamic reorganization of functional brain networks during picture naming. Cortex. 73, 276–288.

Hassan, M., et al., 2016. Identification of interictal epileptic networks from dense-EEG. Brain Topography. 1–17.

Hassan, M., Wendling, F., 2018. Electroencephalography Source Connectivity: Aiming for High Resolution of Brain Networks in Time and Space. IEEE Signal Processing Magazine. 35, 81–96.

Hauk, O., 2004. Keep it simple: a case for using classical minimum norm estimation in the analysis of EEG and MEG data. Neuroimage. 21, 1612–1621.

Hipp, J.F., et al., 2012. Large-scale cortical correlation structure of spontaneous oscillatory activity. Nature neuroscience. 15, 884.

Holmes, C.J., et al., 1998. Enhancement of MR images using registration for signal averaging. Journal of computer assisted tomography. 22, 324–333.

Kabbara, A., et al., 2017. The dynamic functional core network of the human brain at rest. Scientific Reports. 7.

Lachaux, J.-P., et al., 1999. Measuring phase synchrony in brain signals. Human brain mapping. 8, 194–208.

Lachaux, J.-P., et al., 2000. Studying single-trials of phase synchronous activity in the brain. International Journal of Bifurcation and Chaos. 10, 2429–2439.

Liu, Q., et al., 2018. Detecting large-scale brain networks using EEG: impact of electrode density, head modeling and source localization. Frontiers in neuroinformatics. 12, 4.

Michel, C.M., Murray, M.M., 2012. Towards the utilization of EEG as a brain imaging tool. Neuroimage. 61, 371–385.

Nolte, G., et al., 2004. Identifying true brain interaction from EEG data using the imaginary part of coherency. Clinical neurophysiology. 115, 2292–2307.

O’Neill, G.C., et al., 2018. Dynamics of large-scale electrophysiological networks: a technical review. Neuroimage. 180, 559–576.

Palva, J.M., et al., 2018. Ghost interactions in MEG/EEG source space: A note of caution on inter-areal coupling measures. Neuroimage. 173, 632–643.

Rizkallah, J., et al., 2018. Dynamic reshaping of functional brain networks during visual object recognition. Journal of neural engineering. 15, 056022.

Roelfsema, P.R., et al., 1997. Visuomotor integration is associated with zero time-lag synchronization among cortical areas. Nature. 385, 157.

Sakkalis, V., 2011. Review of advanced techniques for the estimation of brain connectivity measured with EEG/MEG. Computers in biology and medicine. 41, 1110–1117.

Schoffelen, J.M., Gross, J., 2009. Source connectivity analysis with MEG and EEG. Human brain mapping. 30, 1857–1865.

Song, J., et al., 2015. EEG source localization: sensor density and head surface coverage. Journal of neuroscience methods. 256, 9–21.

Stam, C.J., Nolte, G., Daffertshofer, A., 2007. Phase lag index: assessment of functional connectivity from multi channel EEG and MEG with diminished bias from common sources. Human brain mapping. 28, 1178–1193.

Tadel, F., et al., 2011. Brainstorm: a user-friendly application for MEG/EEG analysis. Computational intelligence and neuroscience. 2011, 8.

van den Heuvel, M.P., et al., 2016. Associated microscale spine density and macroscale connectivity disruptions in schizophrenia. Biological psychiatry. 80, 293–301.

Van Essen, D.C., et al., 2013. The WU-Minn human connectome project: an overview. Neuroimage. 80, 62–79.

Wang, S.H., et al., 2018. Hyperedge bundling: A practical solution to spurious interactions in MEG/EEG source connectivity analyses. NeuroImage. 173, 610–622.

